# Sugarcane carries relatively larger family of CAMTA transcription factors, active against drought stress

**DOI:** 10.1101/2024.12.26.630382

**Authors:** Vitor Luciano Costa da Silva, Ana Maria Matinez, Melissa Azevedo Vincente, Maqsood Alam, Muhammad Noman, Antonio Chalfun-Junior

## Abstract

As the highly demanding complex genome of the hybrid sugarcane cultivar became publicly available last year (2023), it opened avenues to further study this important crop at molecular level. We are interested in digging the multiple stress responsive transcription factors family, the Calmodulin-Binding Transcription Activator (CAMTA) of sugarcane. This manuscript presents a comprehensive study of *ScCAMTA* family based on the latest sugarcane genome sequence information. Within the 10 gb genome, through HMM model prepared from sorghum CAMTA common domains, we found 48 genes, 46 out of which carry all the CAMTA-associated domains including CG-1, TIG, IQ and Ank. The phylogenetic analysis clustered then into seven classes. Keeping sorghum as reference, we named them as *ScCAMTA1* – *ScCAMTA7*, while each one representing a class having 5-7 copies such as *ScCAMTA1A* – *ScCAMTA1E*, present in each sub-genome (chromosome) within the hybrid sugarcane. In parallel to determining their physico-chemical attributes, gene structure, promoter sequences, protein domains, miRNA targets, protein interaction network, GO, genome collinearity, their expression pattern against drought was determined using the RNAseq data which flaunted that *ScCAMTA7* is highly active under drought. This study furthers the insights into complex sugarcane genome and will assist in developing its drought-tolerant varieties.

## Introduction

The growing global demand for bioenergy and the need for sustainable agricultural practices highlight the importance of sugarcane (Saccharum spp.) as a strategic crop for the development of new energy matrices (Jaiswal et al., 2017). Brazil, as the world’s largest producer, contributed approximately 38% of global sugarcane production, with an estimated output of around 46 million tons of sugar and 35.41 billion liters of ethanol from 8.63 million hectares cultivated in the 2024/2025 harvest season (FAO, 2024; Conab, 2024). In this context, developing more productive and stress-resistant crops represents a key strategy for ensuring a more sustainable future. Detailed insights into the underlying mechanisms of signaling responsible for stress tolerance are promising for helping researchers enhance the natural potential of plant species through biotechnology (Esmaeli et al., 2022; Nascimento et al., 2023; Joshi et al., 2023).

Sugarcane faces significant challenges due to water stress, which can drastically reduce production. Understanding the molecular mechanisms that enable the plant to withstand adverse conditions is essential for developing more resilient varieties (Ferreira et al., 2017). Transcription factors from the CAMTA family (Calmodulin-Binding Transcription Activator) play a key role in calcium-mediated signaling, an essential secondary messenger responding to biotic and abiotic stresses in plants, such as drought, salinity, cold, and pathogenic infections (Galon et al., 2010; Rahman et al., 2016; Wang et al., 2019; Noman et al., 2020). CAMTAs are transcriptional activators that interact with calmodulin (CaM), a calcium-binding protein, to regulate gene expression in response to environmental stimuli. The binding of CaM to calcium enables signal transduction essential for plant adaptation to various stress types. Studies have shown that CAMTAs recognize cis-responsive elements in the promoters of target genes, regulating their expression in response to water stress and other adverse conditions (Yue et al., 2015; Zhou et al., 2022). CAMTAs decode calcium signatures, amplifying Ca2+ signals to produce specific gene expression responses. This process is mediated by the interaction of Ca2+, CaM, and CAMTAs, which amplify calcium signals non-linearly (Liu et al., 2015; Iqbal et al., 2022). Additionally, CAMTAs respond to stress-related hormones, such as abscisic acid, auxin, salicylic acid (SA), and jasmonic acid, modulating gene expression in response to these signals (Yue et al., 2015; Baek et al., 2023). The Ca-CaM-CAMTA pathway is fundamental for plants rapid stress responses, enabling efficient regulation of genes involved in stress tolerance (Yang et al., 2000).

The analysis of the CAMTA family across various plant species has revealed its presence in multicellular organisms and its participation in critical physiological processes. In *Arabidopsis thaliana*, the *AtCAMTA1* and *AtCAMTA3* genes are associated with regulating water stress response and plant immunity (Galon et al., 2010; Nie et al., 2012). Additionally, studies indicate that CAMTAs in soybean (*Glycine max*) play important roles in water stress tolerance, with evidence that their expression is altered under drought conditions (Noman et al., 2019).

Given the complexity of the sugarcane genome, identifying and characterizing CAMTA genes provide valuable insights for genetic improvement aimed at drought resistance. The recent availability of the hybrid sugarcane genome allows for a comprehensive analysis of the *ScCAMTA* family, where 48 genes were identified, 46 of them bearing the domains associated with CAMTA. Phylogenetic analysis grouped these genes into seven distinct classes, suggesting functional diversity to be explored for developing more drought-tolerant varieties.

Transcriptomic analysis revealed that the *ScCAMTA7* gene shows high expression under water stress, indicating its crucial role in regulating sugarcane’s response to adverse conditions. Functional characterization of these genes not only enhances understanding of the molecular mechanisms involved in drought tolerance but also facilitates the development of biotechnological strategies to cultivate more resilient varieties.

In conclusion, the study of CAMTA transcription factors in sugarcane represents a promising approach to addressing the challenges posed by water stress and ensuring the sustainability of agricultural production in Brazil. The intersection of molecular physiology and genetic improvement results in significant innovations to enhance crop resilience in the face of climate change.

## Materials and Methods

### Sequence search and retrieval

As a reference, the sorghum CAMTA protein sequences as well as CAMTA domains such as CG-1, TIG, IQ, and Ank were used against sugarcane genome in Phytozome to search for matching sequences.

In parallel, HMM profiles of the CAMTA gene family (Pfam03859: CG-1; Pfam12796: ankyrin repeats; Pfam01833: TIG domain; and Pfam00612: IQ). An HMM model was created which was used to search for the corresponding sequences in sugarcane. A total of 46 sequences were retrieved. Similarly, the respective sequences in Arabidopsis, rice, maize and sorghum were also downloaded to make a dataset for phylogenetics and subsequent *in silico* analysis.

### Phylogenetic analysis

The CAMTA proteomic sequences of maize, rice, sorghum and Arabidopsis were multiply aligned with *ScCAMTA* dataset, and a phylogenetic tree was generated using MEGA to track their evolutionary relationship. The *ScCAMTA* family was renamed according to phylogenetic dendogram keeping sorghum as reference.

### Chromosomal distribution

The annotated genome sequence file GFF for sugarcane, (retrieved from Phytozome) and related information was input into the TBtools software to visualize the location of all 46 *ScCAMTA* genes along the chromosomes in sugarcane.

### Gene structure

The online tool GSDS was used to visualize the gene structure including the UTR regions and exon-intron assembly using the full-length genomic and CDS sequences of *ScCAMTA* family.

### Physico-chemical properties

The protparam tool at ExPasy determined the physico-chemical properties of *ScCAMTA* from their proteomic sequences.

### Cis-motifs

To check the cis-motifs within the regulatory regions of *ScCAMTA*s, (2kb upstream) 5’ UTR of each gene was resolved using PlantCare database.

### miRNA targets prediction

PsRNATarget database was searched to predict miRNA targets in *ScCAMTA* transcripts.

### Expression analysis in drought based on RNASeq data

RNAseq data in FASTQ format of drought treated sugarcane was retrieved from SRA, processed, and the expression of *ScCAMTA* at various drought stress durations was determined. We used public RNA-seq libraries of the sugarcane hybrid cultivar ROC22, known for its strong resistance to water stress (BioProject: PRJNA776107). Uniform sugarcane seedlings were selected and subjected to a water stress treatment using 20% PEG6000 to simulate drought. Leaf samples were collected at intervals of 0, 4, 8, 16 and 32 hours after the start of treatment, frozen in liquid nitrogen and stored at -80 °C for RNA extraction. The sequencing data was obtained from Li et al. (2022).

The raw reads were subjected to quality check analysis, conducted with FastQC 0.12.1 and high-quality reads were obtained after trimming and filtering the data with Trimmomatic 0.39. Using HISAT2 2.2.1, the raw reads were aligned with the sugarcane hybrid cultivar reference genome (Saccharum officinarum x spontaneum R570 v2.1, https://phytozome-next.jgi.doe.gov/info/SofficinarumxspontaneumR570_v2_1, accessed on 15 June 2024). The StringTie 2.2.3 was used to assemble the reads for comparison and obtain mapped data. The gene expression level was normalized using the Fragments per Kilobase of transcript per Million mapped reads (FPKM) method. All of the downstream analyses were based on high-quality clean data.

## Results

### Phylogenetic analysis

The proteomic sequences of CAMTA families of Arabidopsis, rice, maize, sorghum and sugarcane clustered into 7 clades. In the same clade, *ScCAMTA* are more similar to *ZmCAMTA* and *SbCAMTA*, less similar to wheat, rice, and Arabidopsis. (Naming starts in the sequence of genomes and chromosomes of hybrid sugarcane). *SbCAMTA1, OsCAMTA1, TaCAMTA1A, TaCAMTA1B* and *TaCAMTA1D, ZmCAMTA3, SbCAMTA1, ScCAMTA1D, ScCAMTA1E, ScCAMTA1G, ScCAMTA1C, ScCMATA1B, ScCAMTA1F* and *ScCAMTA1A* clustered together and thus were named as *ScCAMTA1*. Same pattern was followed for the rest of the members. Overall, high similarity for *ScCAMTA* was observed with *SbCAMTA*, which is in agreement with the literature.

**Table 1.**
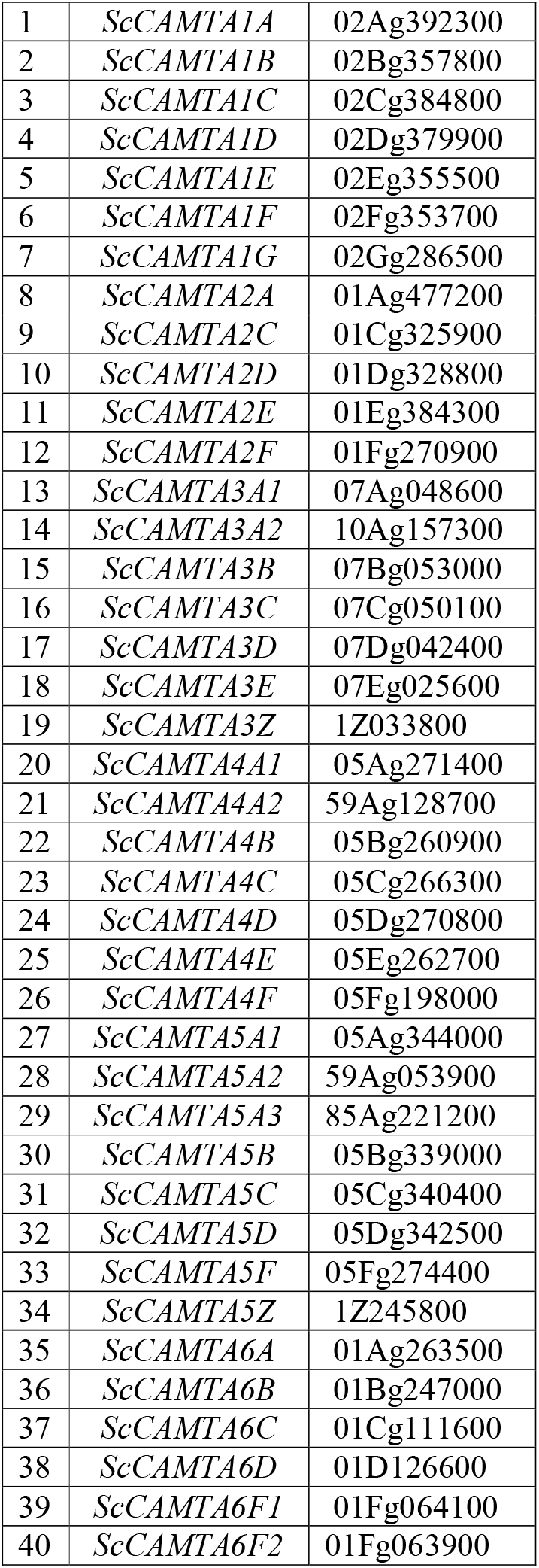
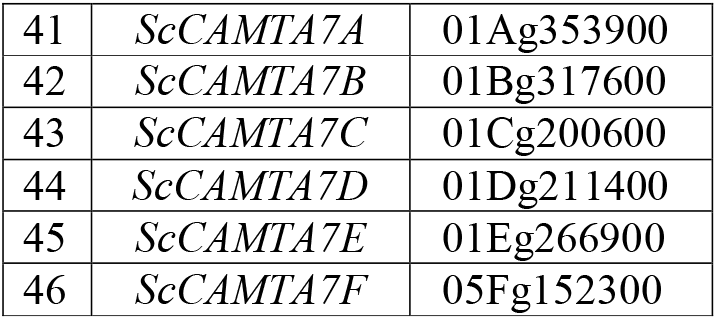
List of 46 members *ScCAMTA* gene family.

### Physico-chemical properties

All ScCAMTA proteins identified are hydrophobic, as GRAVY, with length of amino acid between 841 and 1942 (average: 997), pI value between 5.43 and 8.34 (average: 6.31), Instability index between 37.92 and 51.85 (average: 45.26), and Aliphatic index between 72.02 and 87.81 (average: 76.67).

### Gene Structure

Using the online GSDS tool, the gene structure of all the *ScCAMTA*s was visualized in order to mutually compare their structural diversity. The length of *ScCAMTA* genes lie in the range between 6528bp (ScCAMTA1E) to 42368 bp (ScCAMTA6F2 with an average length of 11374 bp.

### Chromosomal location

The 46 *ScCAMTA* genes are unequally distributed over 27 out of 114 chromosomes and 2 scaffolds of sugarcane R570 hybrid cultivar, as shown in figure 3. The chromosomes 1A, 1C, 1D and 1F has the highest number of *ScCAMTA* genes, i.e., three.

**Figure 1.**
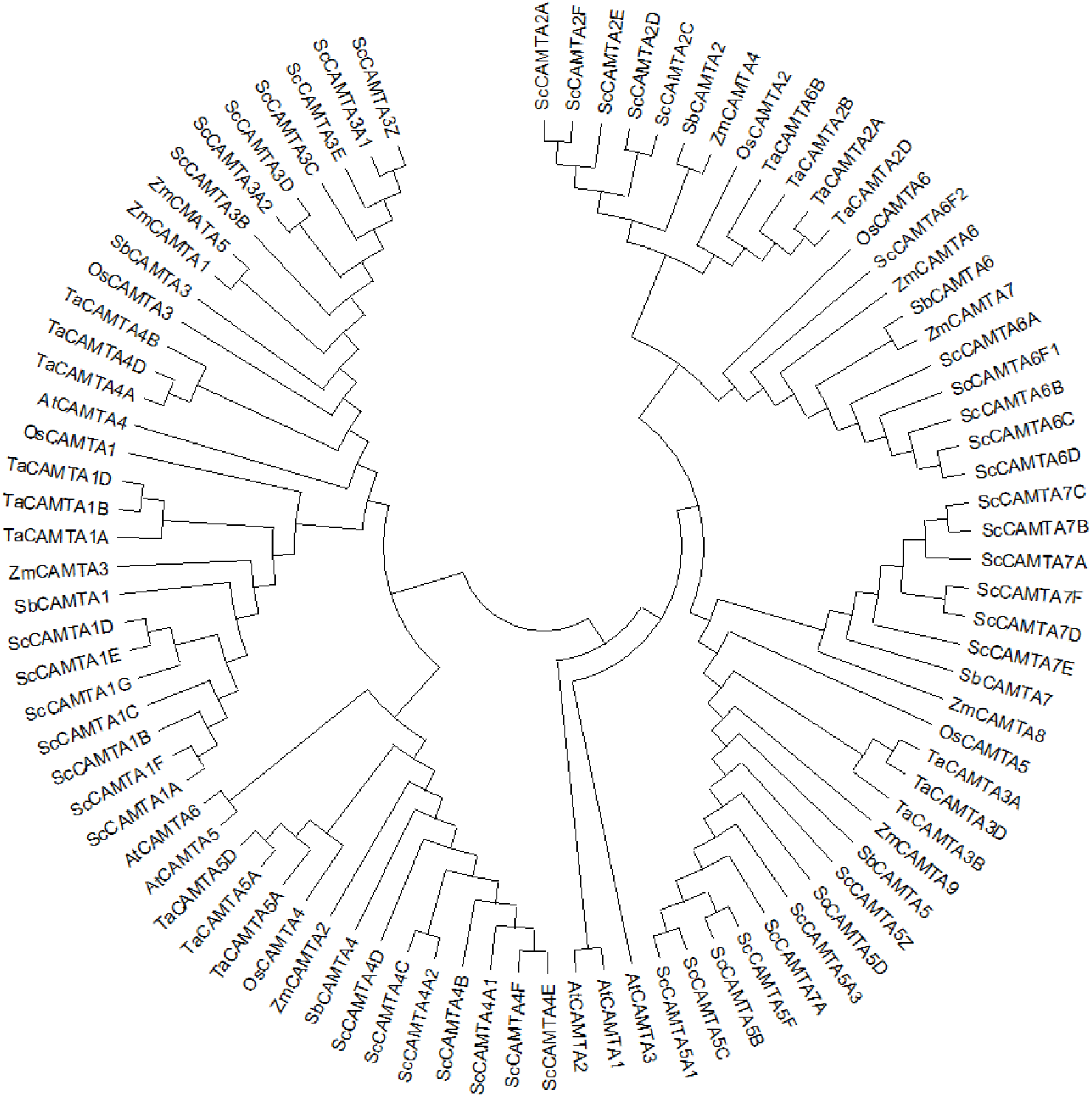
Phylogenetic tree categorizing Sugarcane CAMTAs in 7 distinct clades with respect to Sorghum CAMTAs.

**Figure 2.**
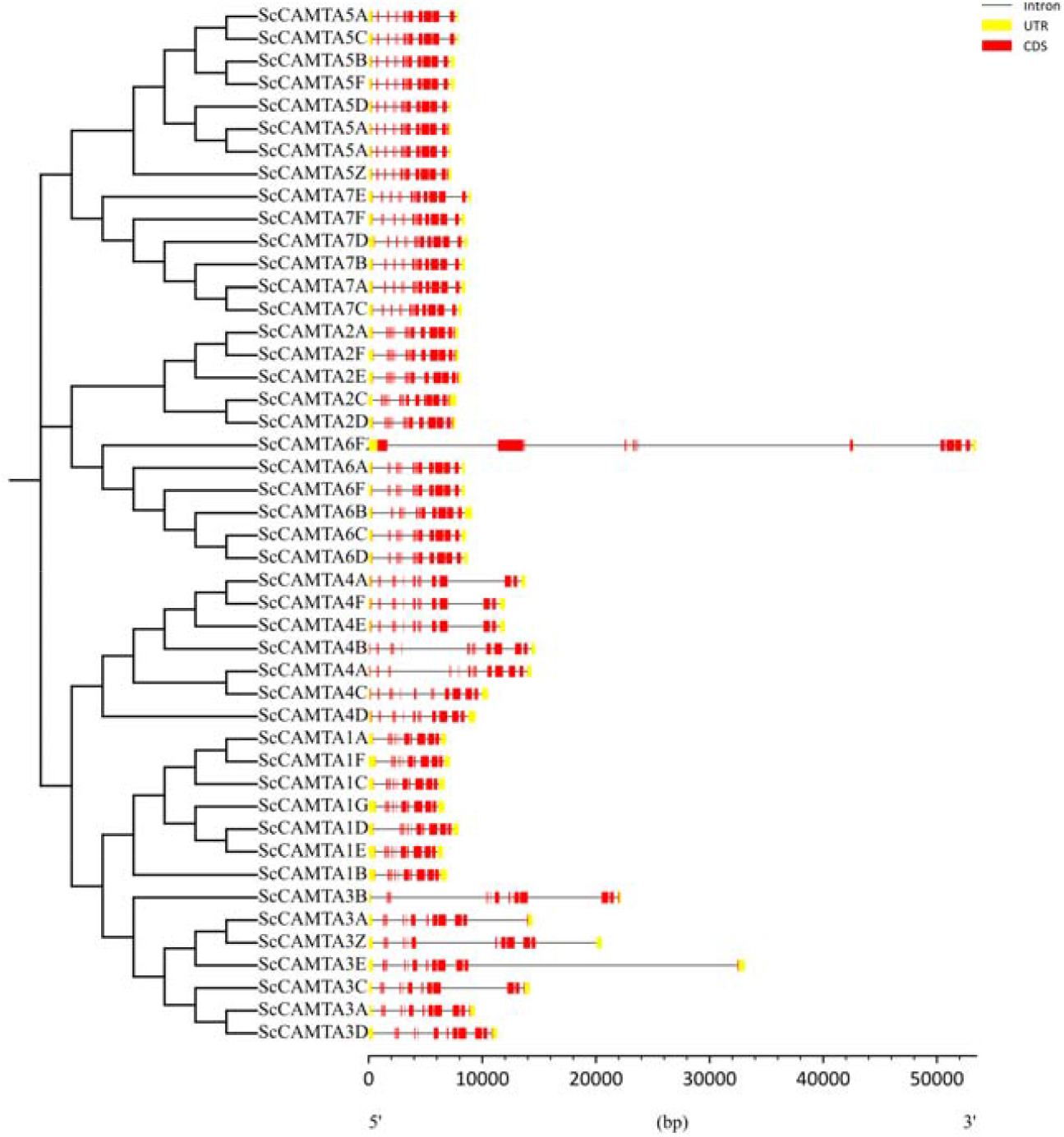
Visualization of the exon-intron assembly of *ScCAMTA* genes with tbtools software.

**Figure 3.**
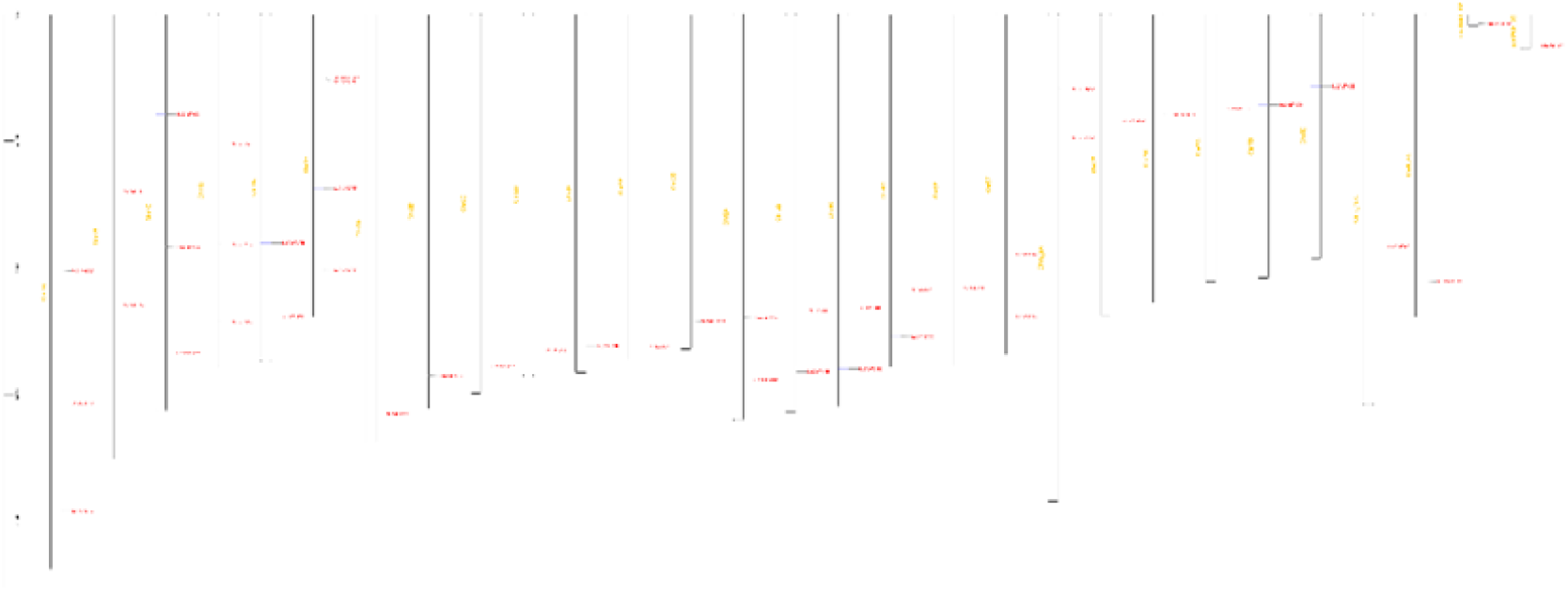
Visualization of the distribution of 46 *ScCAMTA* genes along the chromosomes.

**Figure 5.**
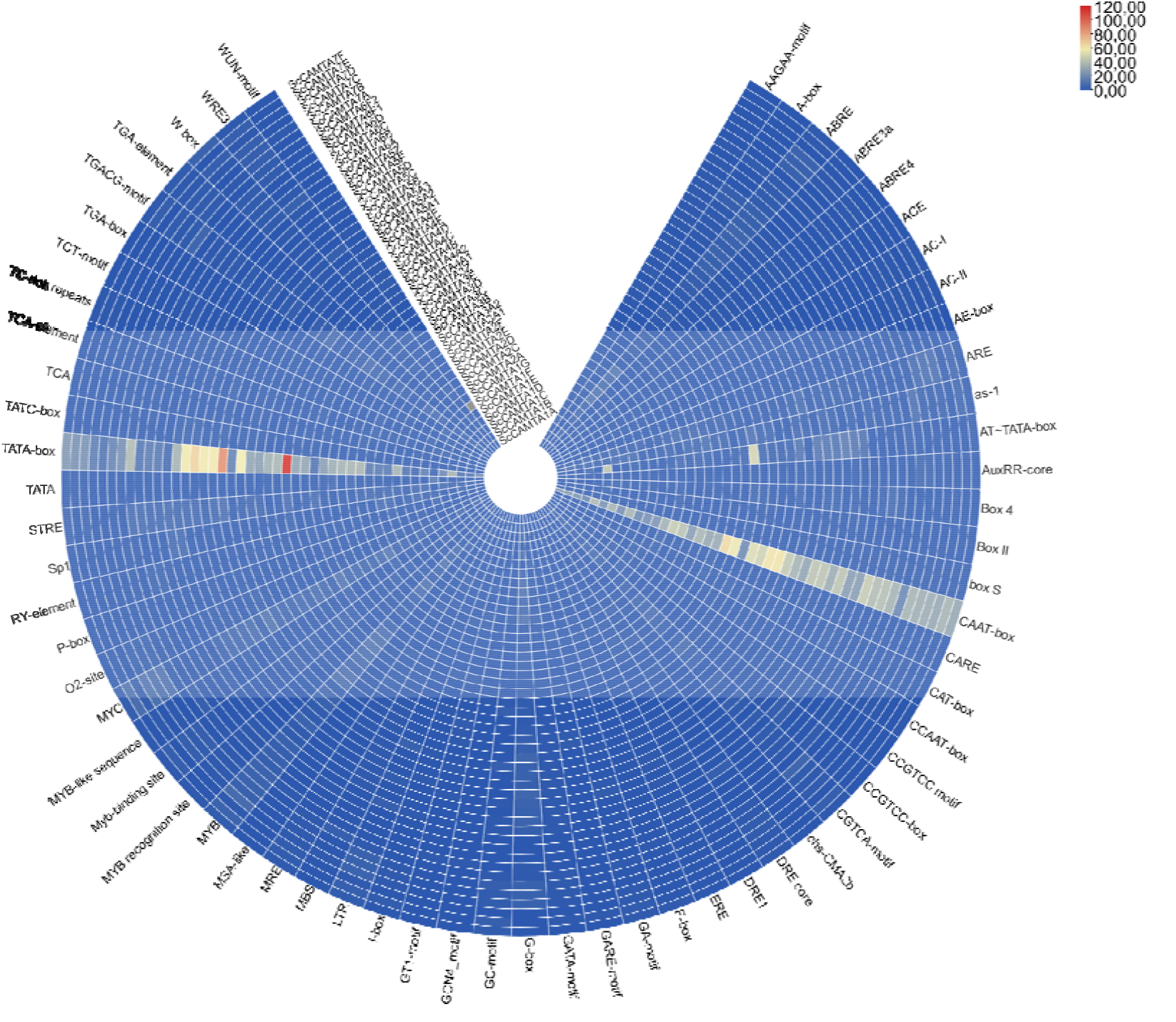
Visualization of cis motifs in the promoter region of *ScCAMTA* genes.

### Cis motifs in the ScCAMTA regulatory sequences

We dissected the regulatory region of ScCAMTAs (-2 kb upstream) which revealed stimulus-specific cis-motifs in their promoters. Overall, there are light (G-box, MRE and AE-box), drought (MBS), salt (MYB), pathogen (TC-rich repeats), wound (WUN, WRE), low temperature (LTR), gibberellin (GARE, P-box), auxin (AUXRR-core) and abscisic acid (ABRE) responsive cis-elements as shown in figure 4. The presence of multiple cis-motifs in *ScCAMTA* genes represents their responsivity to multiple stimuli.

### Synteny analysis

To further insight into the evolutionary relationships of *ScCAMTAs*, a comparative syntenic map of R570 sugarcane hybrid cultivar genome associated with *Sorghum bicolor, Oryza sativa* and *Zea mays* genomes were constructed as shown in figure 6.

**Figure 6.**
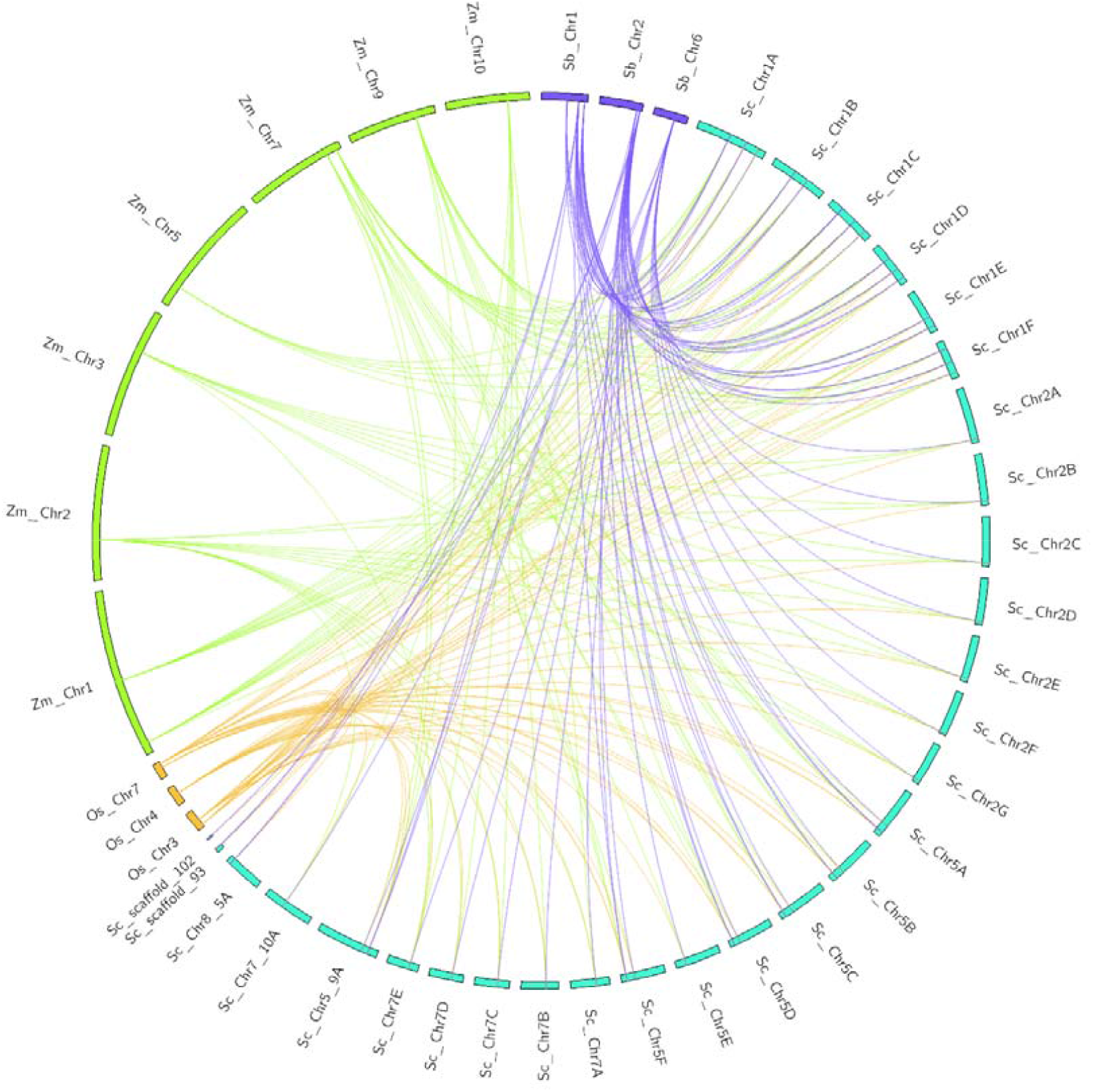
Collinearity of the *CAMTA* genes among rice, maize, sorghum and sugarcane.

### *miRNAs* targeting *ScCAMTA* transcripts

Using the online psRNATarget with E value ≤ 5, a total of 4 unique miRNAs miR444a, miR166, miR396, miR156 were predicted which potentially target the *ScCAMTA* transcripts by inhibiting translation or through cleavage.

### Overview of ROC22 transcriptome

To investigate the transcriptional processes of *ScCAMTA* in ROC22 under drought stress, we analyzed RNA-seq data from this cultivar at 5 time points (0 h, 4 h, 8 h, 16 h, and 32 h)after treatment with PEG6000 (20% w/v), with three biological replicates. Fifteen paired libraries were obtained, totaling 482031418 clean reads. Alignment with the sugarcane reference genome showed an alignment efficiency that ranged from 90.75% to 96.72%. A total of 90,893 genes were obtained when the FPKM was greater than 1.

Principal component analysis (PCA) and cluster heatmap analysis of the transcriptomic data revealed high similarity between the three biological replicates within each treatment. Among treatments, a clear separation was observed between control and stress treatments in ROC22 (Figure 7). Sample cluster heatmap analysis also displayed clusters of similar time points (Figure 8).

**Figure 7.**
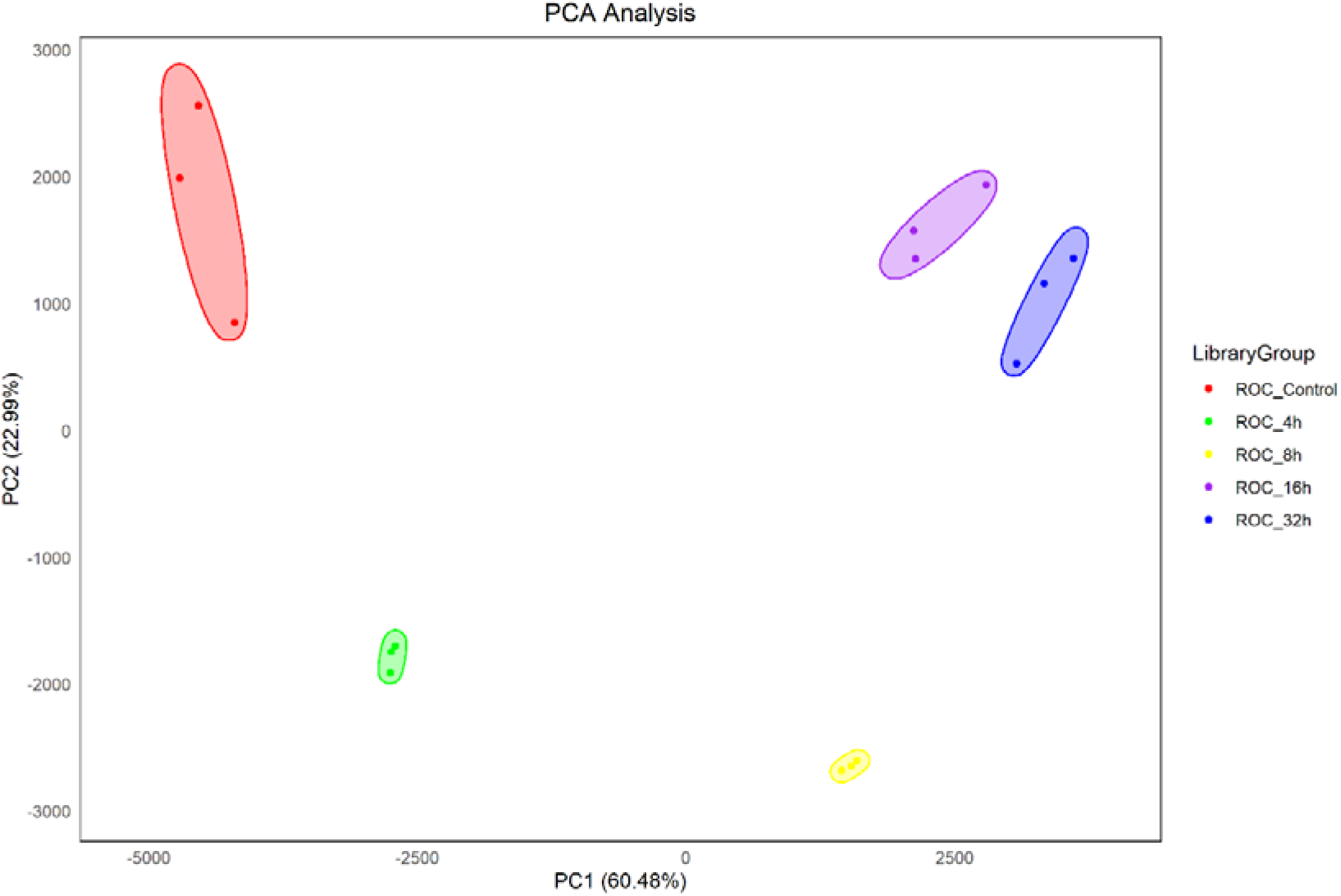
Principal component analysis of the libraries of ROC22 after drought stress treatment along different time points

**Figure 8.**
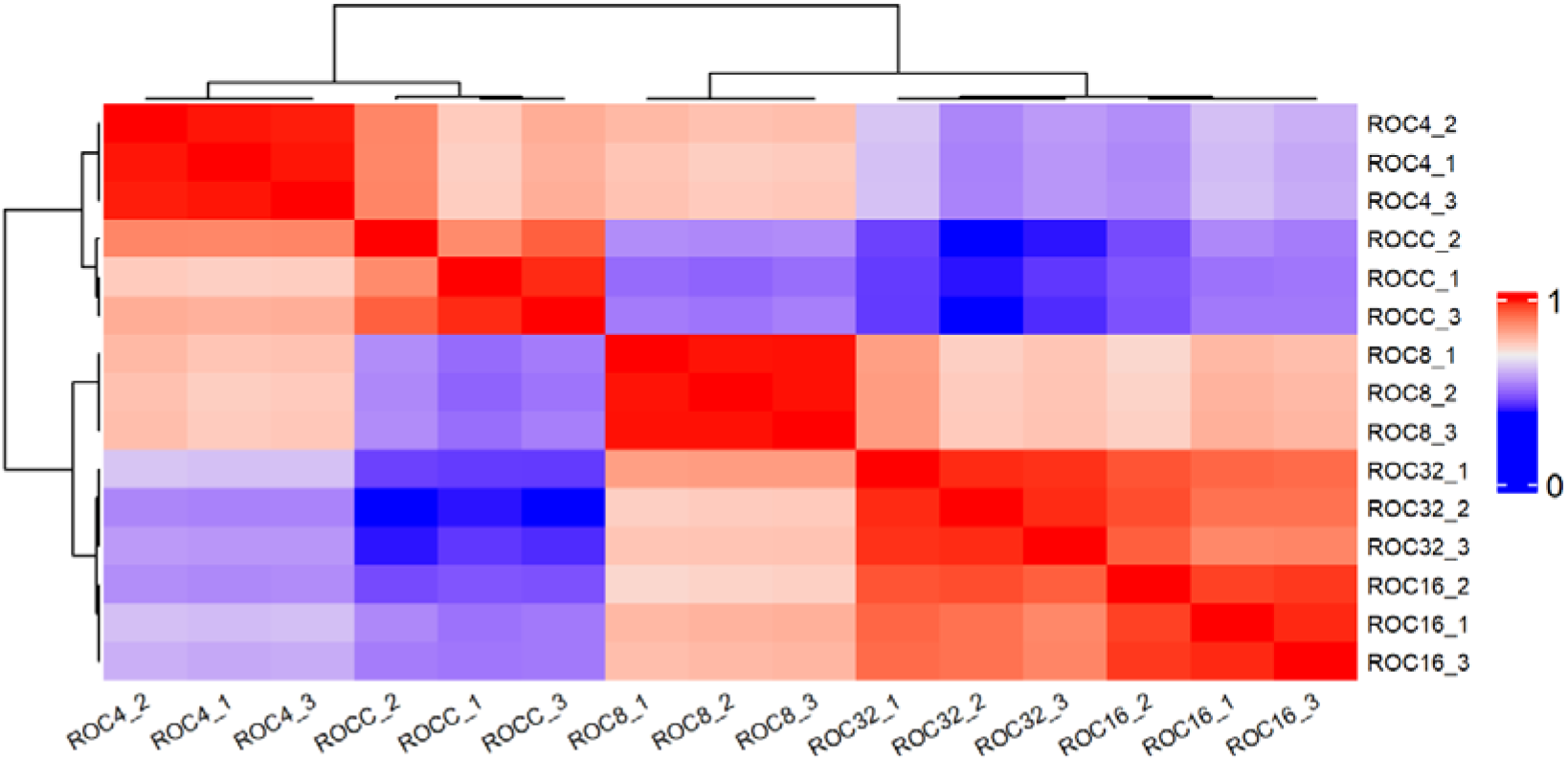
Cluster Heatmap of the libraries of ROC22 after drought stress treatment along different time points

**Figure 9.**
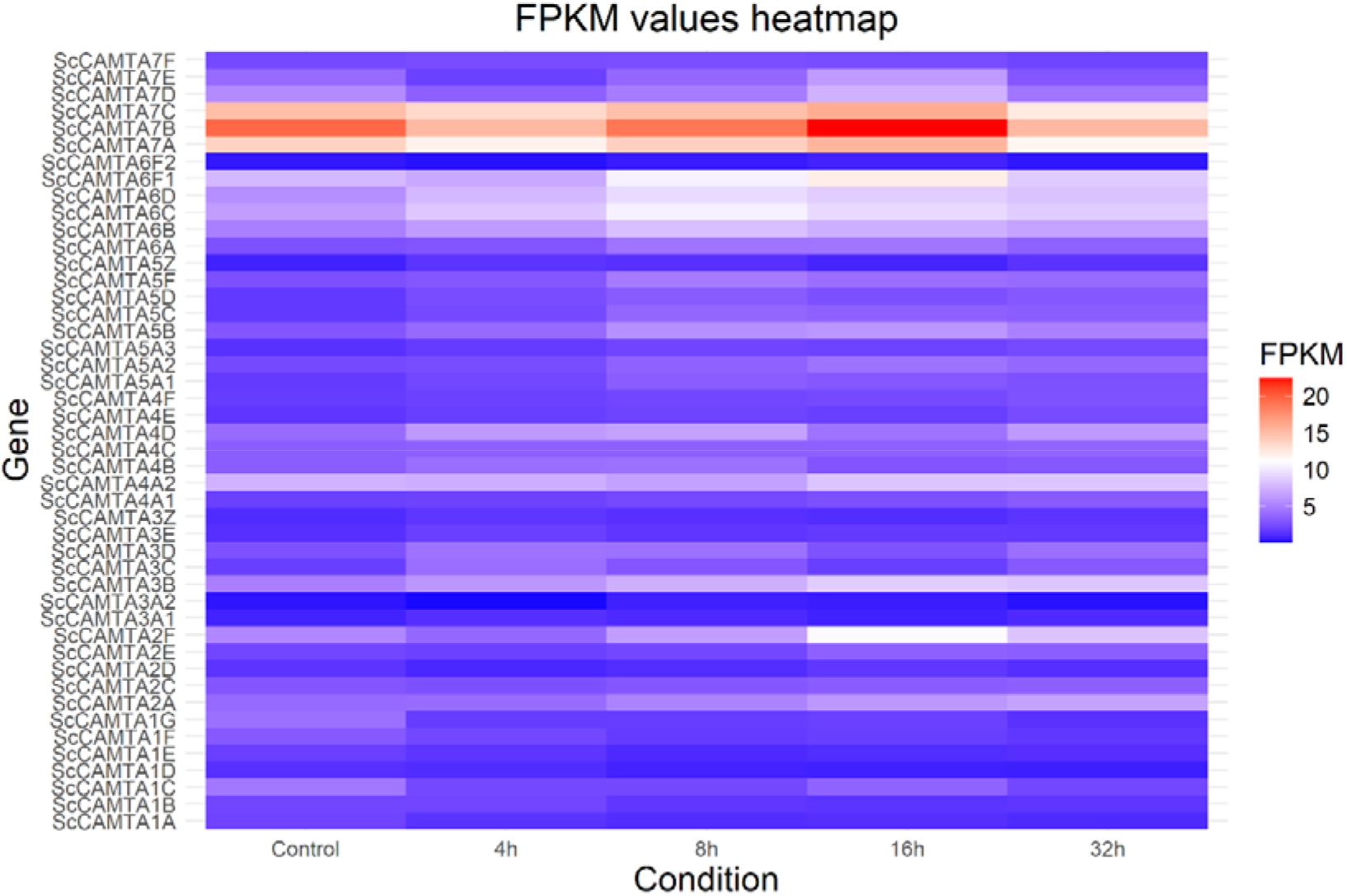
Heatmap of the *ScCAMTA* mean expression between replicates in FPKM after drought stress treatment along different time points

**Figure 10.**
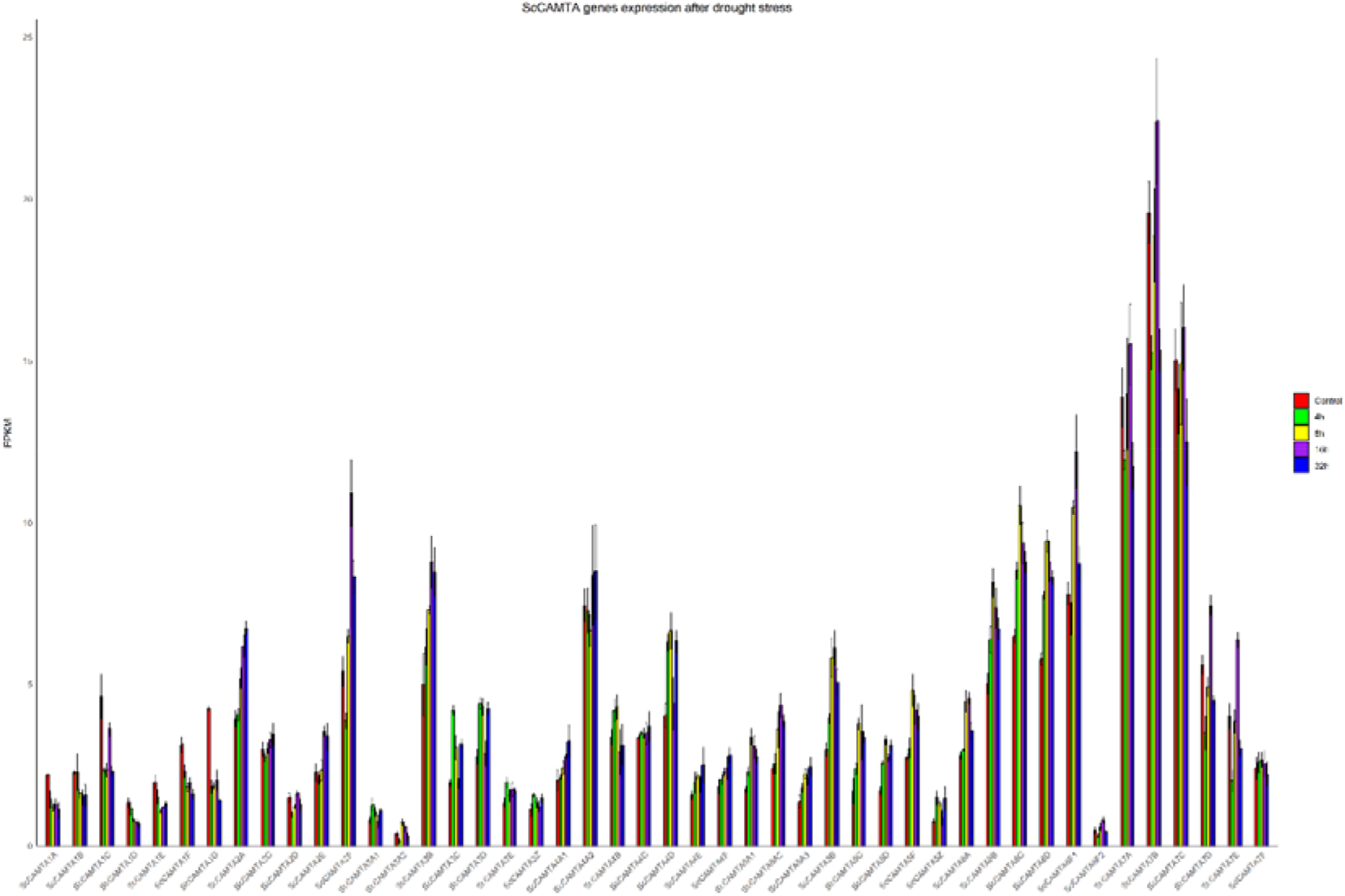
Barplots of the *ScCAMTA* expression in FPKM after drought stress treatment along different time points. Error bars indicates the standard error between the replicates

**Figure 11.**
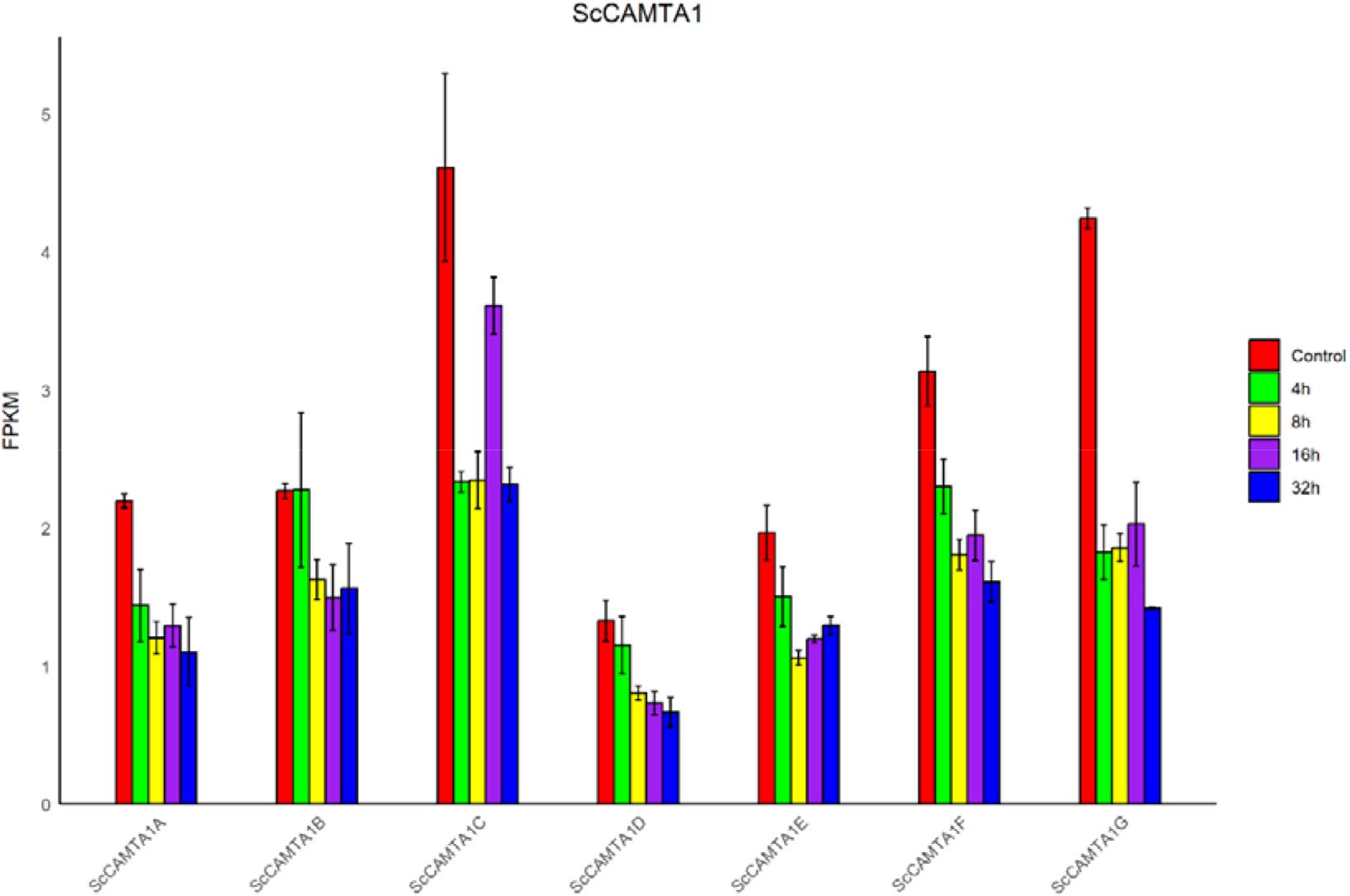
*ScCAMTA1* genes expression in FPKM after drought stress treatment along different time points. Error bars indicate the standard error between the replicates.

**Figure 12.**
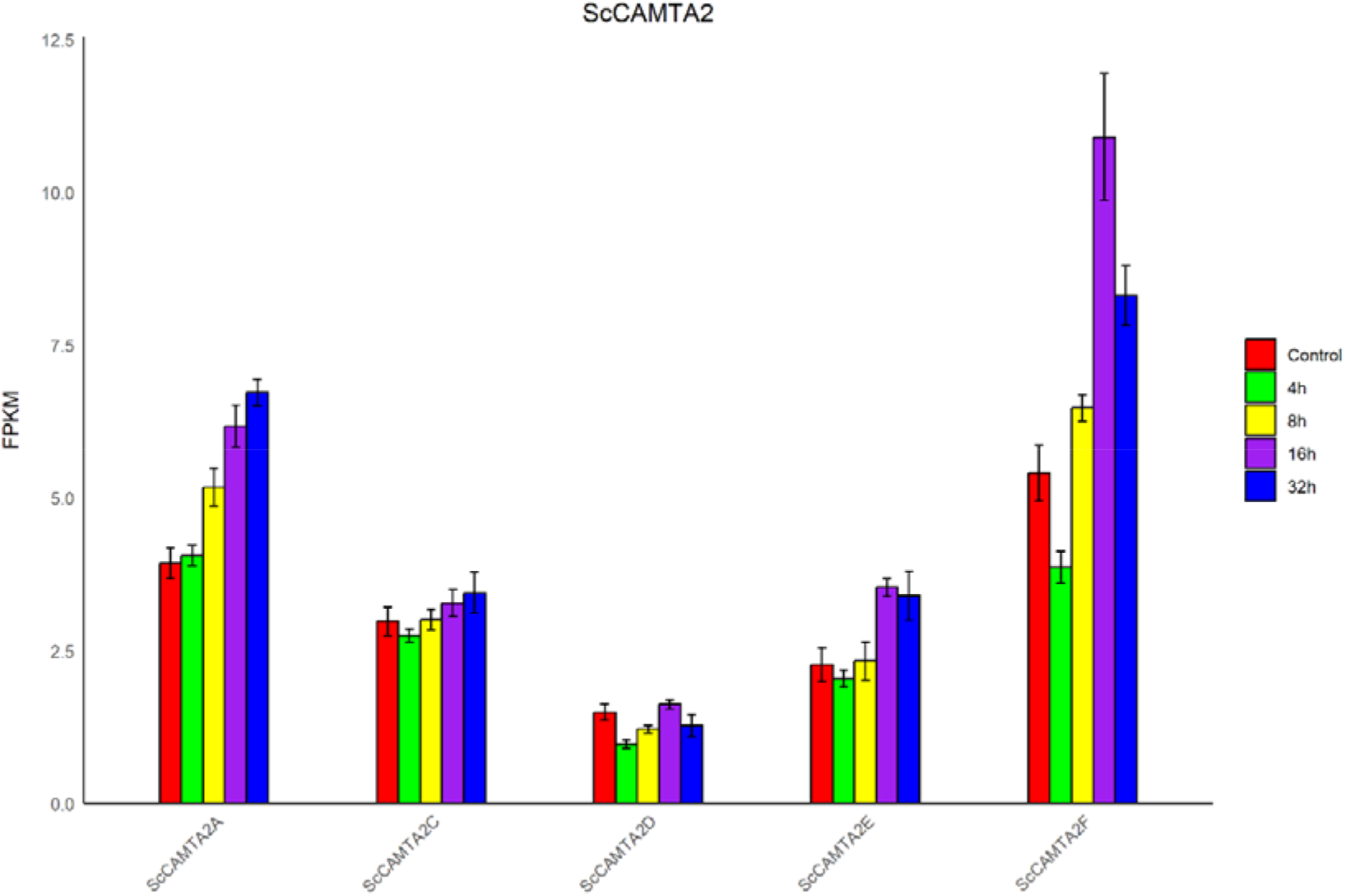
*ScCAMTA2* genes expression in FPKM after drought stress treatment along different time points. Error bars indicate the standard error between the replicates.

**Figure 13.**
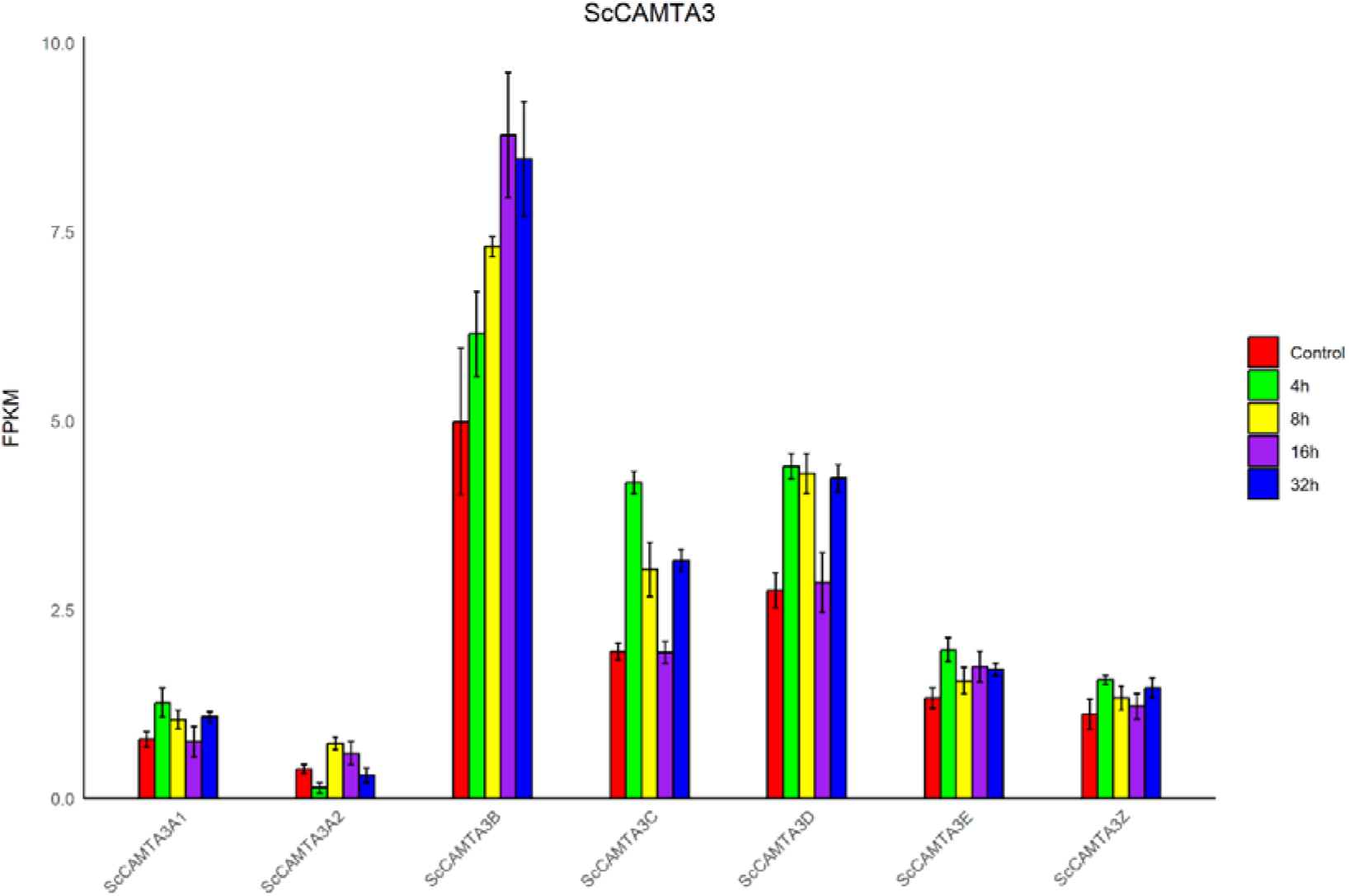
*ScCAMTA3* genes expression in FPKM after drought stress treatment along different time points. Error bars indicate the standard error between the replicates.

**Figure 14.**
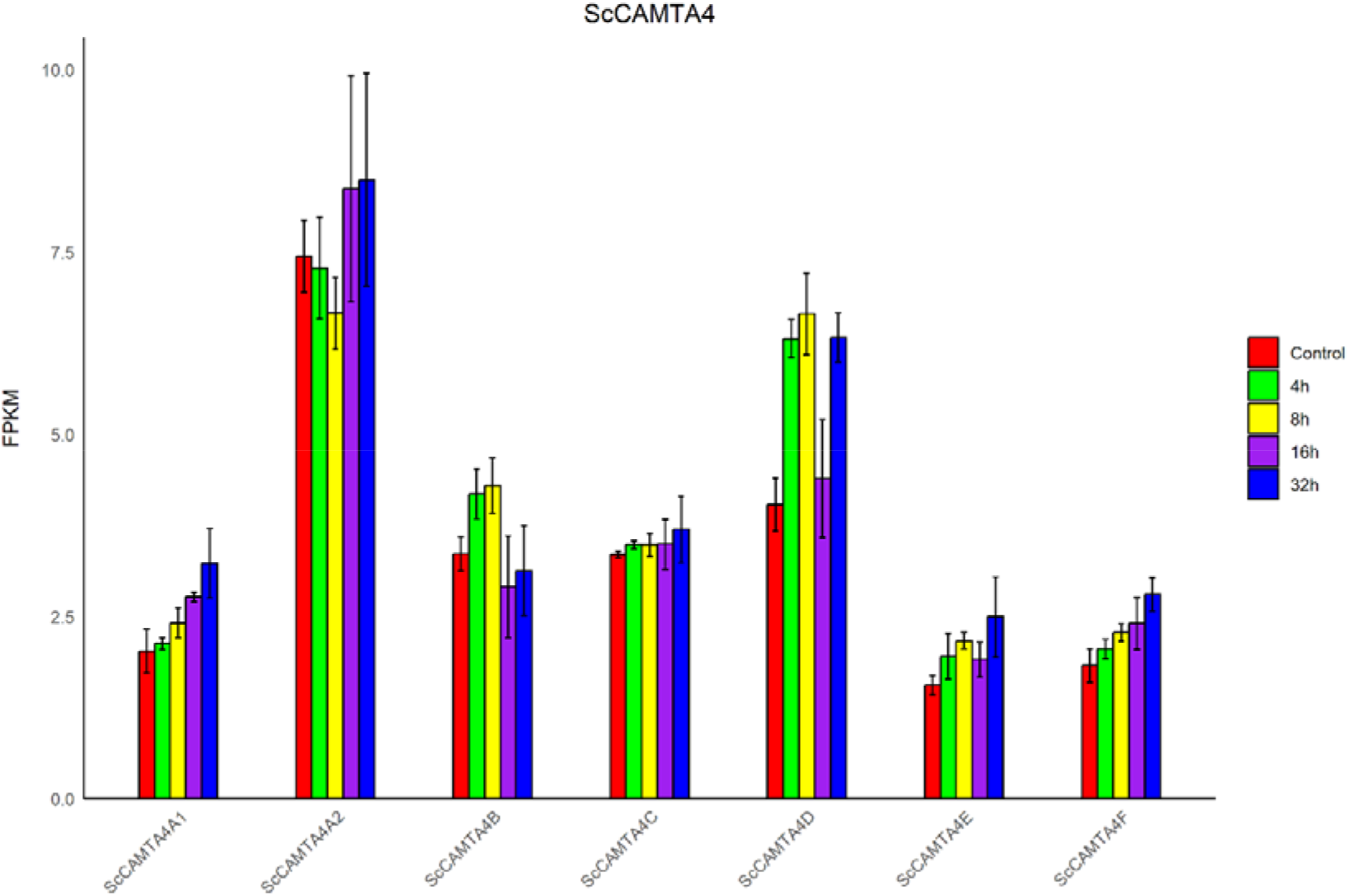
*ScCAMTA4* genes expression in FPKM after drought stress treatment along different time points. Error bars indicate the standard error between the replicates.

**Figure 15.**
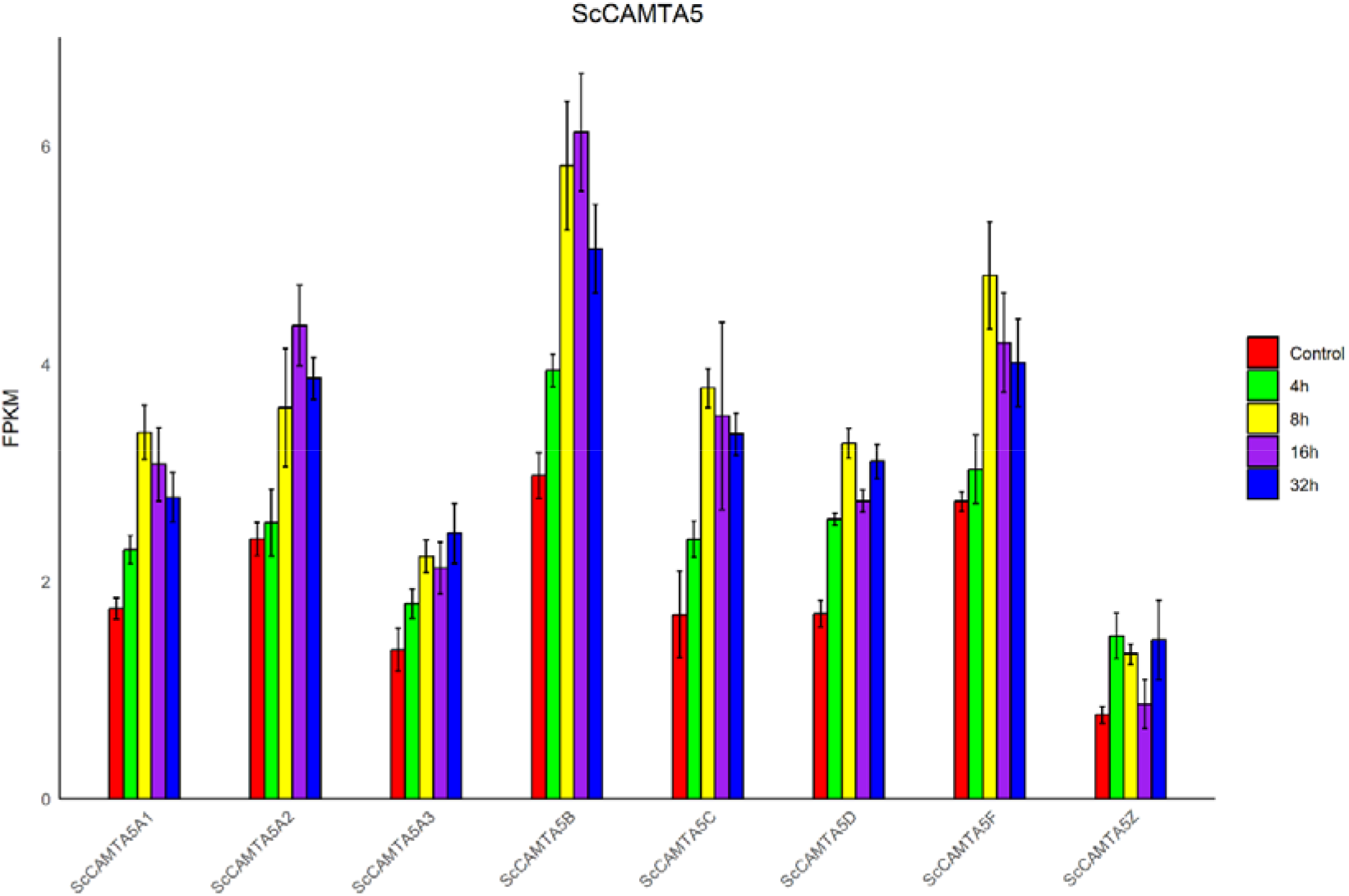
*ScCAMTA5* genes expression in FPKM after drought stress treatment along different time points. Error bars indicate the standard error between the replicates.

**Figure 16.**
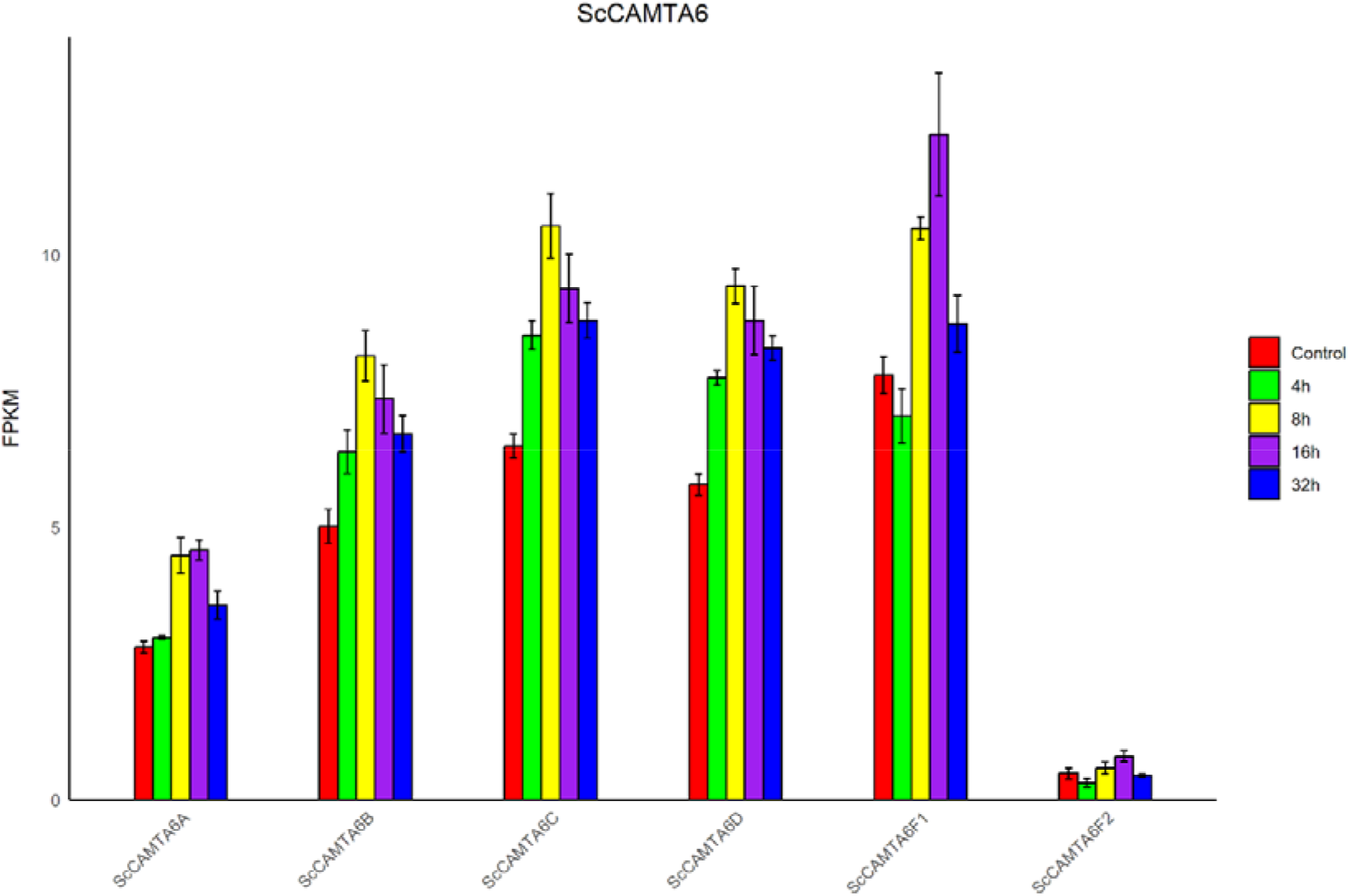
*ScCAMTA6* genes expression in FPKM after drought stress treatment along different time points. Error bars indicate the standard error between the replicates.

**Figure 17.**
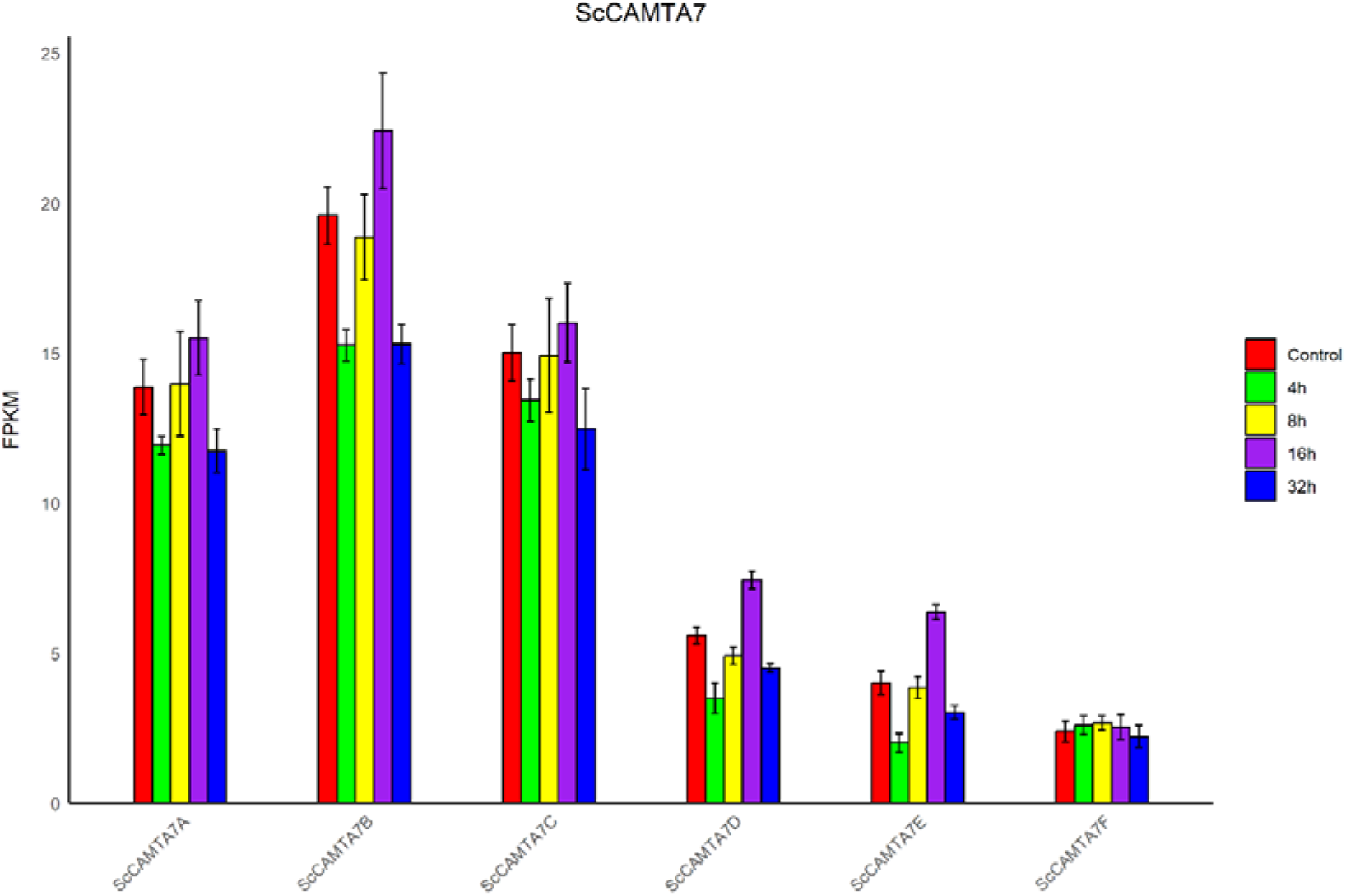
*ScCAMTA7* genes expression in FPKM after drought stress treatment along different time points. Error bars indicate the standard error between the replicates.

### Expression of ScCAMTA genes

Of the 46 genes analyzed, 19 were positively regulated, 6 were negatively regulated, 3 showed neutral behavior and 18 showed neutral behavior indeterminate, demonstrating a distinct regulation between short- and medium-term drought response. The genes with the highest expression, in FPKM, were those of *ScCAMTA7*, homologues of *SbCAMTA7*, indicating its possible role as the main transcription factor in gene regulation of responses to abiotic stresses, via signaling of Ca^2^+. Genes from the *ScCAMTA5* and *ScCAMTA6* clades mostly showed positive regulatory behavior over time, while genes from the *ScCAMTA1* clade showed mostly negative regulation over time.

## References

1. Bordonal, R. D. O., Tenelli, S., da Silva Oliveira, D. M., Chagas, M. F., Cherubin, M. R., Weiler, D. A., … & Carvalho, J. L. N. (2024). Carbon savings from sugarcane straw-derived bioenergy: Insights from a life cycle perspective including soil carbon changes. Science of the Total Environment, 947, 174670. 10.1016/j.scitotenv.2024.174670

2. Jaiswal, D., De Souza, A. P., Larsen, S., LeBauer, D. S., Miguez, F. E., Sparovek, G., … & Long, S. P. (2017). Brazilian sugarcane ethanol as an expandable green alternative to crude oil use. Nature Climate Change, 7(11), 788–792. 10.1038/nclimate3410

3. FAO. 2024. World Food and Agriculture – Statistical Yearbook 2024.

4. Conab, C. D. A. (2024). Acompanhamento da safra Brasileira de cana-de-açúcar.–v. 12.

5. Esmaeili, N., Shen, G., & Zhang, H. (2022). Genetic manipulation for abiotic stress resistance traits in crops. Frontiers in Plant Science, 13. 10.3389/fpls.2022.1011985.

6. Nascimento, F., Rocha, A., Soares, J., Mascarenhas, M., Ferreira, M., Lino, L., De Souza Ramos, A., Diniz, L., Mendes, T., Ferreira, C., Santos-Serejo, J., & Amorim, E. (2023). Gene Editing for Plant Resistance to Abiotic Factors: A Systematic Review. Plants, 12. 10.3390/plants12020305.

7. Joshi, A., Yang, S., Song, H., Min, J., & Lee, J. (2023). Genetic Databases and Gene Editing Tools for Enhancing Crop Resistance against Abiotic Stress. Biology, 12. 10.3390/biology12111400.

8. Ferreira, T., Tsunada, M., Bassi, D., Araújo, P., Mattiello, L., Guidelli, G., Righetto, G., Gonçalves, V., Lakshmanan, P., & Menossi, M. (2017). Sugarcane Water Stress Tolerance Mechanisms and Its Implications on Developing Biotechnology Solutions. Frontiers in Plant Science, 8. 10.3389/fpls.2017.01077.

9. Galon, Y., Aloni, R., Nachmias, D. et al. Calmodulin-binding transcription activator 1 mediates auxin signaling and responds to stresses in Arabidopsis. Planta 232, 165–178 (2010). 10.1007/s00425-010-1153-6

10. Rahman, H., Yang, J., Xu, Y. P., Munyampundu, J. P., & Cai, X. Z. (2016). Phylogeny of plant CAMTAs and role of AtCAMTAs in nonhost resistance to Xanthomonas oryzae pv. oryzae. Frontiers in plant science, 7, 177. 10.3389/fpls.2016.00177

11. Rahman, H., Xu, Y. P., Zhang, X. R., & Cai, X. Z. (2016). Brassica napus genome possesses extraordinary high number of CAMTA genes and CAMTA3 contributes to PAMP triggered immunity and resistance to Sclerotinia sclerotiorum. Frontiers in plant science, 7, 581. 10.3389/fpls.2016.00581

12. Wang, Y., Wei, F., Zhou, H. et al. TaCAMTA4, a Calmodulin-Interacting Protein, Involved in Defense Response of Wheat to Puccinia triticina. Sci Rep 9, 641 (2019). 10.1038/s41598-018-36385-1

13. Noman, M., Aysha, J., Ketehouli, T., Yang, J., Du, L., Wang, F., & Li, H. (2020). Calmodulin binding transcription activators: An interplay between calcium signalling and plant stress tolerance. Journal of plant physiology, 256, 153327 . 10.1016/j.jplph.2020.153327.

14. Zhou, Q., Zhao, M., Xing, F., Mao, G., Wang, Y., Dai, Y., Niu, M., & Yuan, H. (2022). Identification and Expression Analysis of CAMTA Genes in Tea Plant Reveal Their Complex Regulatory Role in Stress Responses. Frontiers in Plant Science, 13. 10.3389/fpls.2022.910768.

15. Yue, R., Lu, C., Sun, T., Peng, T., Han, X., Qi, J., Yan, S., & Tie, S. (2015). Identification and expression profiling analysis of calmodulin-binding transcription activator genes in maize (Zea mays L.) under abiotic and biotic stresses. Frontiers in Plant Science, 6. 10.3389/fpls.2015.00576.

16. Liu, J., Whalley, H., & Knight, M. (2015). Combining modelling and experimental approaches to explain how calcium signatures are decoded by calmodulin□binding transcription activators (CAMTAs) to produce specific gene expression responses. The New Phytologist, 208, 174–187. 10.1111/nph.13428.

17. Iqbal, Z., Memon, A., Ahmad, A., & Iqbal, M. (2022). Calcium Mediated Cold Acclimation in Plants: Underlying Signaling and Molecular Mechanisms. Frontiers in Plant Science, 13. 10.3389/fpls.2022.855559.

18. Baek, D., Cho, H., Cha, Y., Jin, B., Lee, S., Park, M., Chun, H., & Kim, M. (2023). Soybean Calmodulin-Binding Transcription Activators, GmCAMTA2 and GmCAMTA8, Coordinate the Circadian Regulation of Developmental Processes and Drought Stress Responses. International Journal of Molecular Sciences, 24. 10.3390/ijms241411477.

19. Yang, T., & Poovaiah, B. W. (2000). An early ethylene up-regulated gene encoding a calmodulin-binding protein involved in plant senescence and death. Journal of Biological Chemistry, 275(49), 38467–38473. 10.1074/jbc.M003566200

20. Nie, H., Zhao, C., Wu, G., Wu, Y., Chen, Y., & Tang, D. (2012). SR1, a calmodulin-binding transcription factor, modulates plant defense and ethylene-induced senescence by directly regulating NDR1 and EIN3. Plant physiology, 158(4), 1847–1859.

